## Supplemental Material for "Evolutionarily conserved chaperone-mediated proteasomal degradation of a disease-linked aspartoacylase variant"

*Fig. S1 (ASPA aggregates do not inhibit yeast growth and do not localize to JUNQ or IPOD*): *p2*

*Fig. S2 (Prediction of DnaK binding sites in ASPA):* *p3*

*Fig. S3 (Hsp110 is primarily required for degradation of newly synthesized ASPA):* *p4*

*Fig. S4 (Cdc48 promotes degradation of insoluble ASPA):* *p5*

*Fig. S5 (ASPA degradation is ubiquitin-dependent):* *p6*

*Fig. S6 (Knock-down of HSPA4 results in increased levels of insoluble ASPA):* *p7*

*Fig. S7 (Hsp110-mediated ASPA degradation):* *p8*

*Supplemental references:* *p9*

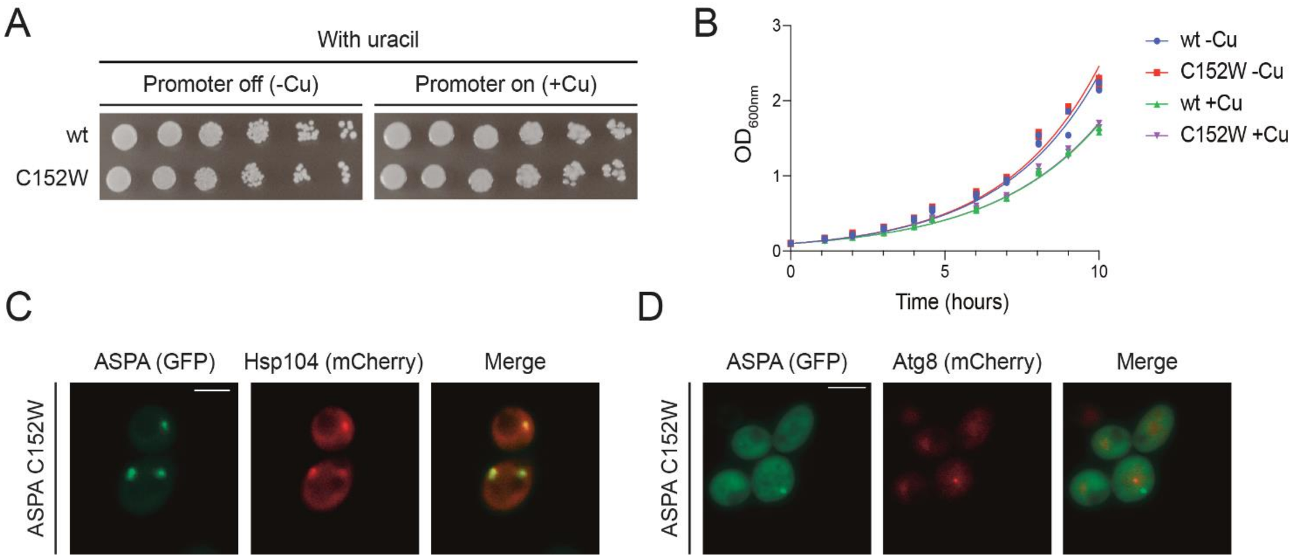


**Figure S1** *ASPA aggregates do not inhibit yeast growth and do not localize to JUNQ or IPOD*. (A) Wild-type yeast cells expressing either wild-type ASPA or the C152W variant grown on solid medium containing uracil and without or with copper as indicated. (B) Wild-type yeast expressing the indicated constructs were grown in liquid medium containing uracil without or with copper as indicated, and the $OD_{600nm}$ was measured at the indicated time points. Growth was compared using Tukey's multiple comparisons test on three independent cultures for each condition. There was no significant difference between the doubling time of wild-type ASPA or the C152W variant in absence nor in presence of copper. The doubling times in absence and presence of copper were significantly different (adjusted p-value < 0.0001, n=3) possibly caused by copper itself or the high expression level of an exogenous protein. (C) Representative fluorescence microscopy images of wild-type yeast expressing GFP-ASPA C152W and Hsp104-mCherry. ASPA aggregates consistently co-localized with Hsp104, as would be expected for both the JUNQ and the IPOD (Kaganovich et al., 2008). Scale bar is 4 µm. (D) Representative fluorescence microscopy images of wild-type yeast expressing GFP-ASPA C152W and mCherry-Atg8. Only rarely did we observe co-localization between ASPA aggregates and Atg8, indicating that aggregated ASPA C152W does not accumulate in the IPOD (Kaganovich et al., 2008). Scale bar is 4 µm.

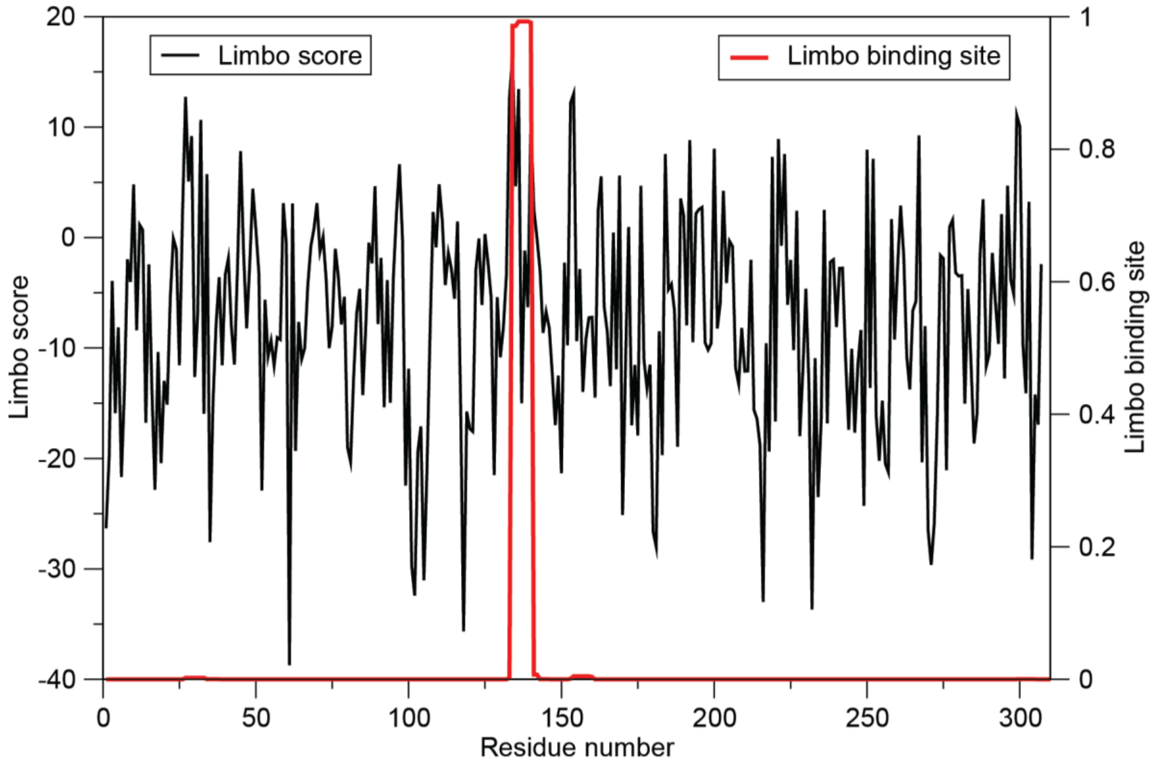


**Figure S2** *Prediction of DnaK binding sites in ASPA.* We used Limbo (Van Durme et al., 2009) to predict putative DnaK (Hsp70) binding sites in the ASPA sequence, and show both (black) the raw binding scores (over seven residue peptides) and (red) a "Boltzmann" probability to highlight the most likely binding sites.

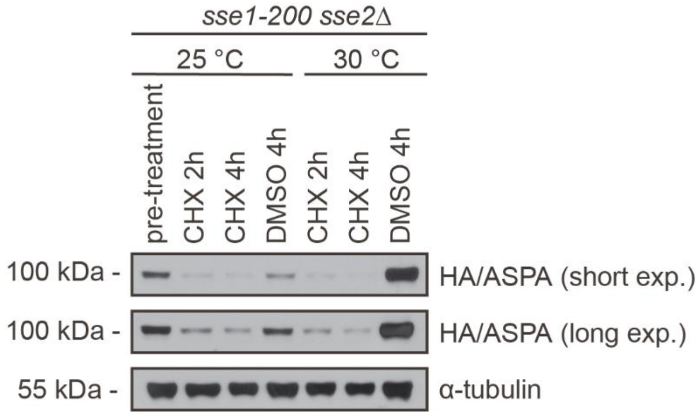


**Figure S3** *Hsp110 is primarily required for degradation of newly synthesized ASPA.* The *sse1-200 sse2Δ* strain expressing ASPA C152W was incubated at 25 °C until exponential phase. The culture was then split and cells were incubated at the permissive temperature (25 °C) or restrictive temperature (30 °C) with the translation inhibitor cycloheximide (CHX) or as a control the solvent DMSO. Cells were harvested at the indicated time points, followed by protein extraction and Western blotting to examine the steady state of ASPA C152W. Tubulin served as a loading control.

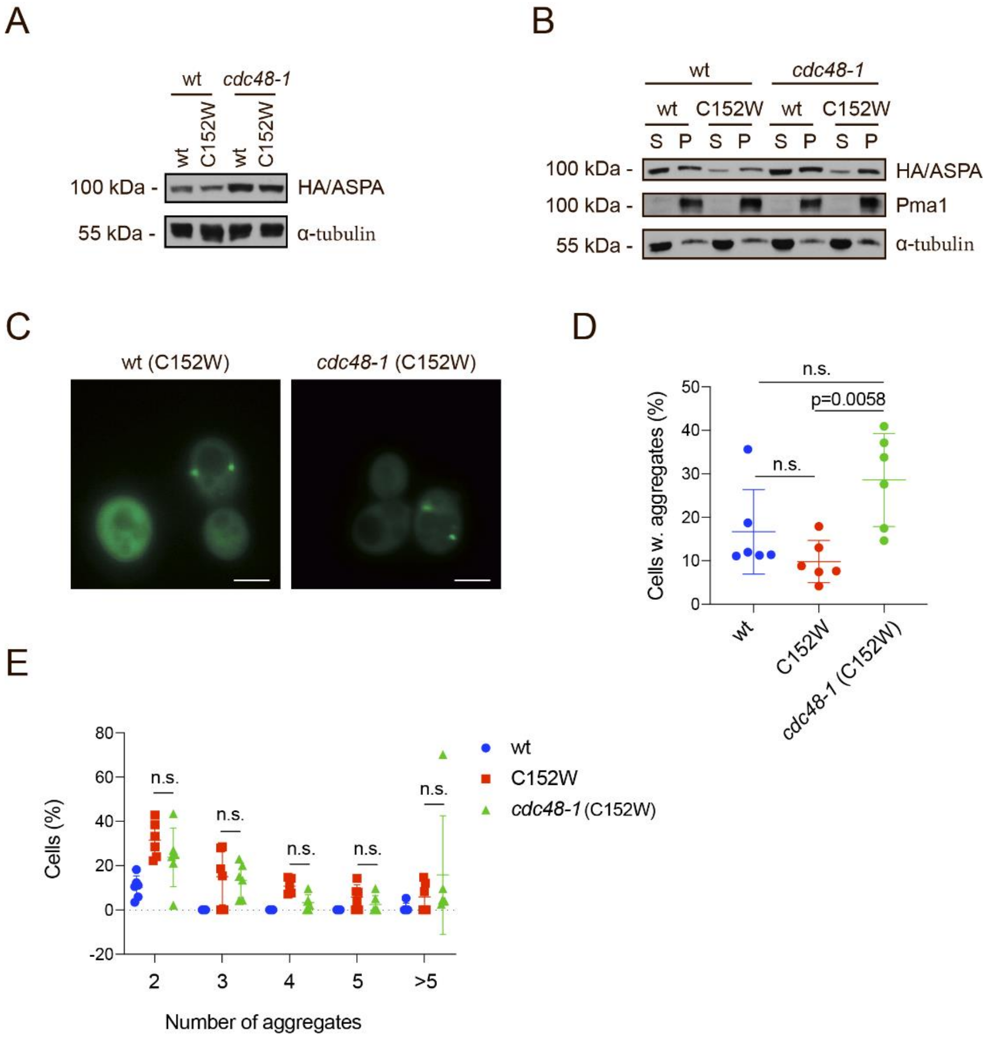


**Figure S4** *Cdc48 promotes degradation of insoluble ASPA.* (A) Western blot of ASPA protein levels in wild-type and *cdc48-1* yeast strains expressing the indicated constructs cultured with 0.1 mM $CuSO_4$. Tubulin served as a loading control. (B) Western blot showing the levels of soluble (S) and insoluble (P) ASPA in wild-type and *cdc48-1* yeast strains expressing the indicated constructs in presence of 0.1 mM $CuSO_4$. Tubulin and Pma1 served as controls for the soluble and insoluble fractions, respectively. (C) Representative fluorescence microscopy images of GFP-ASPA C152W in a wild-type or *cdc48-1* strain. Scale bars are 4 µm. (D) Cells expressing GFP and containing aggregates were counted using fluorescence microscopy of wild-type (wt, C152W) and *cdc48-1* yeast strains expressing the indicated constructs. Data points indicate frequencies from independent experiments (minimum of 127 GFP-expressing cells, n=6). The frequencies of cells containing aggregates were compared using a one-way ANOVA and Tukey's multiple comparisons test in GraphPad Prism. Means and standard deviations are shown. (E) Cells containing aggregates and the number of aggregates were counted by fluorescence microscopy of wild-type (wt, C152W) and *cdc48-1* yeast cells expressing the shown constructs. The frequencies of each observation in separate experiments are shown as data points (minimum of 127 GFP-expressing cells, n=6). The aggregate patterns were compared using a two-way ANOVA and Tukey's multiple comparisons test. Means and standard deviations are shown.

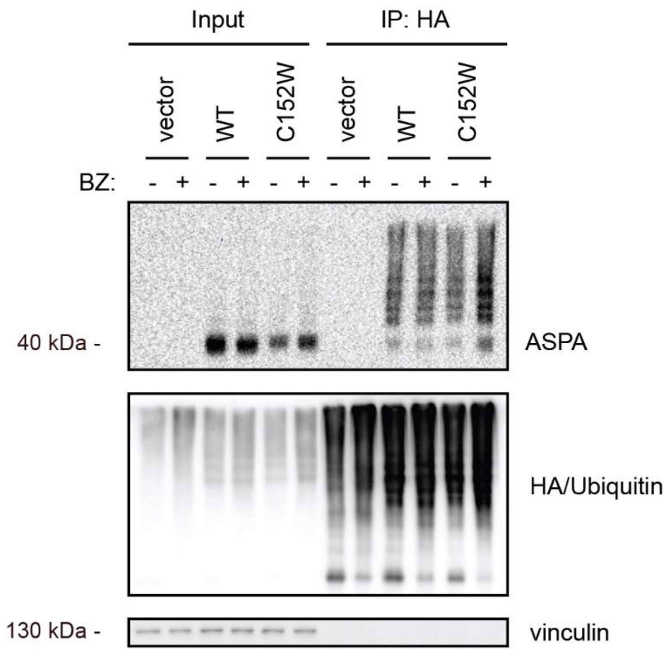


**Figure S5** *ASPA degradation is ubiquitin-dependent.* U2OS cells transiently transfected to express HA-ubiquitin and a vector control, wild-type ASPA or C152W were treated with bortezomib (BZ, +) or as a control DMSO (-) followed by denaturing HA immunoprecipitation. Vinculin serves as a loading control. Note that the level of ubiquitinated ASPA C152W increases following treatment with bortezomib.

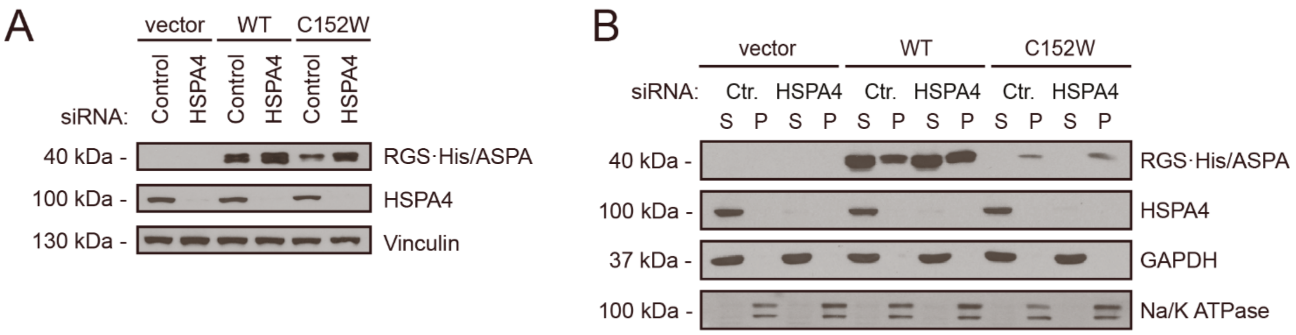


**Figure S6** *Knock-down of HSPA4 results in increased levels of insoluble ASPA.* (A) Knock-down of *HSPA4* in U2OS cells transiently transfected with the indicated constructs results in stabilization of ASPA. Successful knock-down was confirmed by blotting for HSPA4. Vinculin served as a loading control. (B) Cells were treated as in A, followed by differential centrifugation to separate soluble and insoluble protein into supernatant (S) and pellet (P) fractions. Successful knock-down was confirmed by blotting for HSPA4. GAPDH and the Na/K ATPase served as controls for soluble and insoluble proteins, respectively. Note that ASPA C152W is exclusively found in the insoluble fraction.

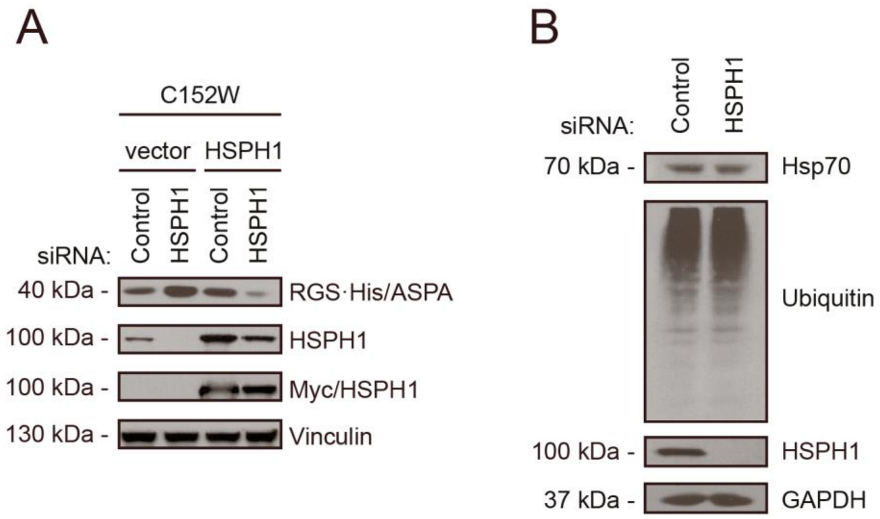


**Figure S7** *Hsp110-mediated ASPA degradation.* (A) U2OS cells transiently transfected to express ASPA C152W and HSPH1 or a vector control with and without *HSPH1* knock-down. Note that expression of episomal siRNA-resistant myc-tagged HSPH1 restores degradation of ASPA C152W upon knock-down of endogenous *HSPH1*. Vinculin served as a loading control. (B) Knock-down of HSPH1 expression does not lead to detectable changes in the level of Hsp70 or ubiquitin-protein conjugates. GAPDH served as a loading control.
